# Mathematical model of RNA-directed DNA methylation predicts tuning of negative feedbacks required for stable maintenance

**DOI:** 10.1101/2023.12.05.570286

**Authors:** Renee Dale, Rebecca A. Mosher

**Affiliations:** Donald Danforth Plant Sciences Center, St. Louis, MO, 63132; School of Plant Sciences, University of Arizona, Tucson, AZ 85721-0036

**Author notes:** Author for correspondence: RD.

**Keywords:** siRNA, DNA Methylation, Argonaute, RNA Pol IV, RdDM

## Abstract

RNA-directed DNA Methylation (RdDM) is a plant-specific *de novo* methylation pathway that is responsible for maintenance of asymmetric methylation (CHH, where H=A, T, or G) in euchromatin. Loci with CHH methylation are transcriptionally silent and produce 24-nucleotide (nt) short interfering (si) RNAs. These siRNAs direct additional CHH methylation to the locus, thereby maintaining methylation states through DNA replication. To understand the necessary conditions to produce stable CHH methylation, we developed a stochastic mathematical model of RdDM. The model describes DNA target search of DNA or RNA by siRNAs derived from CHH-methylated loci. When the siRNA (bound by an Argonaute protein) finds the matching locus, the model uses the dwell time of the matched complex to determine the degree of CHH reinforcing methylation. Reinforcing methylation occurs either throughout the cell cycle (steady reinforcement), or immediately following replication (bursty reinforcement). Each simulation occurs over 10 cell cycles, and for 7 simulation replicates. We use nonparametric statistics to compare initial and final CHH methylation distributions to determine whether the simulation conditions produce stable maintenance. We apply this method to the low CHH methylation case, wherein the median is only 8%, and many loci have less than 8% methylation. The resulting model predicts that siRNA production must be linearly proportional to CHH methylation levels at each locus, that bursty reinforcement produces more stable systems, and that slightly higher levels of siRNA production are required for DNA target search, compared to RNA target search. Unlike CG methylation, which typically exhibits bi-modality, with loci having either 100% or 0% methylation, CHH methylation putatively exists at a range of methylation fractions. Our model predicts that careful tuning of the negative feedbacks in the system are required to balance the positive feedback loop of increasing CHH methylation and increasing siRNA production, enabling stable maintenance of a range of CHH methylation across replication events.

## Introduction

In eukaryotic genomes, cytosines are methylated to influence the behavior of DNA, and loss of this methylation has profound effects on gene expression and genome stability [1,2]. DNA methylation in animals is primarily on cytosines that are immediately 5’ to a guanine (a “CG” site), whereas DNA methylation in plants occurs on cytosines regardless of the surrounding sequence context. Distinct mechanisms maintain CG, CHG, and CHH methylation (where H=A, T, or C) in plant genomes. Most DNA methylation occurs in large blocks of heterochromatin, the highly condensed and gene-poor regions of the genome. This methylation persists following DNA replication because the signals recruiting methyltransferases are encoded in histone modifications, which are equally partitioned to the two daughter strands. The 50% reduction in modified histones is sufficient to restore DNA methylation levels before the next round of DNA replication. In contrast, CHH methylation in euchromatin, the portion of the genome containing most protein-coding genes, is maintained by RNA-directed DNA Methylation (RdDM) [3].

RdDM employs a complex mechanism whereby 24-nucleotide (nt) small interfering (si)RNAs are synthesized from methylated loci, processed and bound by Argonaute proteins (AGO) in the cytoplasm, and reimported to the nucleus before targeting DNA methylation based on the sequence information encoded by the siRNA [4,5]. Binding of the AGO:siRNA complex to a target locus recruits a *de novo* methyltransferase to place additional CHH methylation. Many of the components necessary for RdDM have been identified via forward or reverse genetic screens and biochemical characterization of these components has led to a complex schematic model of this mechanism. RdDM begins when RNA Pol IV produces short non-coding transcripts [6,7]. Pol IV then backtracks along DNA, releasing the 3’ end of the transcript and passing this to RNA-DEPENDENT RNA POLYMERASE 2, which uses it as a template for synthesis of a complementary strand [8]. The short double-stranded RNAs produced by these polymerases are substrates for DICER-LIKE 3, which trims them to produce 24-nt siRNA duplexes [9,10]. These siRNA duplexes are exported to the cytoplasm, possibly via TREX/THO complex [11], where they are bound by an Argonaute (AGO) protein. The 24-nt siRNA produced in RdDM are bound by Argonautes in the AGO4-clade [12] and siRNA binding triggers nuclear localization of AGO4:siRNA complexes [13]. In the nucleus, the AGO:siRNA complex uses the sequence of its siRNA to identify complementary nucleic acids – probably non-coding transcripts produced by RNA Pol V, or potentially single stranded DNA liberated during Pol V transcription [14–17]. Localization of the AGO:siRNA complex at chromatin recruits DOMAINS REARRANGED METHYLTRANSFERASE (DRM) which methylates cytosines regardless of their sequence context [18–21]. The DNA methylation triggered by DRM causes methylation of Histone H3 Lysine 9 (H3K9me), a mark that recruits Pol IV for further siRNA production [20,22,23]. DRM has preference for double-stranded DNA, but preferentially methylates only one of the two strands [19]. However, bidirectional siRNA production and Pol V transcription results in methylation of both DNA strands at a target locus [24]. DNA methylation is therefore distributed to both daughter strands following DNA replication; similarly, methylated histones are randomly distributed to daughter strands, resulting in a strong feedback loop to maintain DNA methylation through cell divisions [20,25].

Despite this detailed molecular model, there are a number of unanswered questions regarding the mechanism of RdDM. For example, although non-coding RNA produced by Pol V is generally assumed to be the target of AGO:siRNA complexes, zero-distance crosslinking localizes AGO4 to the DNA, suggesting that AGO:siRNA complexes directly bind to DNA [16]. The carboxy terminal domain of RNA Pol V also contains numerous AGO hook motifs, which bind AGO proteins in a sequence-independent manner [26]; these motifs are required for RdDM, but sequence-independent binding is not part of the canonical model. In addition to unanswered questions regarding the mechanism, we also have no quantitative understanding of RdDM and the parameters that enable maintenance of methylation and siRNA production through many cell divisions. It is particularly notable that sites of RdDM differ in both the amount of siRNA produced and level of methylation, yet these different levels are consistent between individuals, indicating a system that is stable at a range of parameter values [27–29]. The median levels of RdDM methylation vary greatly, often around 20% methylation, and occasionally as low as 8% [30].

In addition to diagrammatic models that describe the simple relationship between components, biological processes can also be described by mathematical models that incorporate dynamic and quantitative interactions between the components. Mathematical models allow researchers to determine the quantitative parameters of the biological process, and also to test characteristics of the system and discover new relationships or components [31]. *De novo* and maintenance DNA methylation have been modeled using stochastic models and coupled rate equations [32–40]. However, these models focus on CG methylation, whose maintenance is fundamentally different from RdDM maintenance of CHH methylation. After semi-conservative DNA replication, CG sites become hemi-methylated, and hemi-methylated sites are directly recognized by DNMT1-type methyltransferases. In contrast, at CHH sites one duplex remains methylated, while the other is unmethylated until acted upon by RdDM [3]. Mathematical modeling has also been applied to RNA silencing mechanisms [41–43]; however, these models have been limited to post-transcriptional silencing of mRNA transcripts in the cytoplasm, rather than small RNA-mediated modification of DNA or chromatin.

Here, we produce the first quantitative mathematical model for RdDM and we investigate the parameter values necessary to produce stable methylation across multiple cell divisions. We focus on the necessary conditions to produce a stable system under the low methylation case, where the median CHH methylation is only 8%. We demonstrate that the relationship between methylation and siRNA production is linear and that RdDM is likely limited to a discrete portion of the cell cycle for intermediate methylation states to exist. We also demonstrate that both AGO4-RNA and AGO4-DNA associations are feasible in our model.

## Methods

### Model development

The RdDM maintenance model was developed in Matlab 2021a (The MathWorks Inc. (2021. MATLAB version: 5.32.0 (R2021a), Natick, Massachusetts: The MathWorks Inc. https://www.mathworks.com), as described in the supplemental methods. Briefly, 1000 unique loci were modeled and a randomly assigned a CHH methylation level drawn from the distribution −*log*(1 − (1 − *exp*(−*μ*)) * *U* / *μ*, where *μ* is the mean methylation fraction, and *U* is uniformly-distributed noise, 1 − .9 * *Unif*(0,1). This distribution was chosen as it can take a variety of forms (left or right skewed, symmetric, and uniformly distributed). We do not consider variation between initial conditions as significant due to the high sample size (1000). A pool of siRNAs are generated from the loci based on the methylation present at the locus; because the loci are unique, each siRNA matches only a single locus. SiRNAs compete for association with a limited number of AGO proteins and these AGO:siRNA complexes then search for appropriate loci. If an AGO:siRNA complex interacts with a non-matching locus, it will slide along the locus or disassociate and potentially interact with a new locus.

Movement of siRNA:AGO complexes between loci was modeled based on established parameters describing facilitated diffusion across RNA or DNA strands, such as short-distance diffusion (*i*.*e*. sliding, hopping), longer jumps and strand transfer, and dissociation from the locus (exit) [44–47] (**Table 1, Supplemental Text 1**). If an AGO:siRNA complex associates with its matching locus, the time-to-disassociation is determined by the RNA-RNA or RNA-DNA dissociation constant (*k*_*D*_). The total “dwell time” (sum of duration of association of all matching AGO:siRNA complexes) at a given locus determines the amount of methylation placed during the cell cycle. The level of methylation decreases by half at the start of each cell cycle due to new DNA synthesis. Similarly, the AGO:siRNA population is randomly reduced by half and new siRNAs are synthesized. A description of the model algorithm is provided as **Supplemental Text 1**.

**Table 1.** Model parameters and sources. The complete siRNA sequence consists of the seed sequence with 3’ and 5’ supplementary sequences. *pb: photobleaching limit when determined by FRET.

| Parameter | Value | Value in model | Source |
| --- | --- | --- | --- |
| AGO:siRNA affinity | 10<sup>-30</sup> nM (AGO2:mRNA)<br>10<sup>-80</sup> nM (AGO2:miRNA)<br>7.2 nM (AGO2:siRNA) | Ignored; Recycling of AGO:siRNA pool done through replication events | [48–52] |
| AGO limiting | Likely limiting (RISC) | Typically 0.8 AGO : 1 siRNA; not significant influence (confounded with siRNA production level) | [53] |
| AGO level | Coupled to siRNA abundance | Coupled to siRNA abundance | [53,54] |
| AGO degradation | Unstable | AGO is limited in proportion to siRNA abundance | [54] |
| AGO:siRNA degradation | Very stable (0.0004/s degradation, AGO2) | Ignored; Recycling of AGO:siRNA pool done through replication events | [52] |
| siRNA production per unit CHH methylation | Unknown | Several relationships tested |  |
| siRNA production rate | Unknown | Internally estimated using Median Total Dwell Time |  |
| AGO:siRNA affinity to RNA strand | $k_{on}$: 3.9 x 10<sup>8</sup> nM/s<br>$k_{off}$: 0.0036 (<pb*)<br>10pm to 10 nM (RISC)<br>0.18 nM $k_D$ (AGO2:siRNA) | $k_{off}$: 0.02 (less than seed + 3' supplementary) | [44,52,55] |
| AGO:siRNA affinity to DNA strand | $k_{on}$: 1*10<sup>9</sup> nM/s<br>$k_{off}$: 0.41 /s (<pb) | $k_{off}$ is 0.03 for DNA (near detection limit but higher than RNA) | [44] |
| AGO searching RNA rate | 50% to move left or right<br>$k_{on}$ 3.9*10<sup>8</sup> /Ms | 50% to move left or right<br>Shuttling every 1s (calculated - 3x faster than DNA) | [44,47] |
| AGO searching DNA rate | 50% L/R<br>1 x 10<sup>9</sup>/Ms $k_{on}$ | 50% L/R<br>Shuttling every 0.3s (calculated - 3x slower than RNA) | [44,47] |
| AGO searching: jumping | 10% | 10% stochastic coinflip | [47] |
| RNA off-targets | 50% | 50% chance of a transcript coming from an siRNA-producing locus |  |
| DNA off-targets | 90% | 90% of DNA does not produce siRNAs |  |
| Dwell time to produce 1% increase in CHH methylation |  | Internally estimated using Median Total Dwell Time |  |

### Model simulation workflow

The model was run across a range of parameters, including the siRNA production level (50-700 times methylation fraction); the methylation saturation point for siRNA production; steady or bursty siRNA production; for different relationships between siRNA production and methylation level (linear, sigmoid, Hill function); and for AGO:siRNA binding to RNA or DNA targets. With a linear relationship, we calculate the number of siRNA produced as methylation level multiplied by the siRNA production level, such that for 10% methylation and an siRNA production level of 100, 10 siRNAs are produced from that locus. When modeling a linear relationship with a saturation point, we take the minimum result of either the linear relationship or the saturation point multiplied by the siRNA production level. For the Hill function relationship, we calculated the number of siRNA produced using 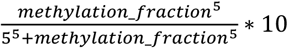. For the sigmoidal Hill function relationship, we used the following form,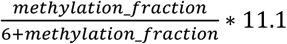. Each simulation was run for 10 generations with initial loci CHH methylation levels drawn from the same starting methylation distribution. Each simulation condition had 7 simulation replicates with 1000 randomly drawn methylation fractions. During the first burn-in generation the dwell time (sum of duration of association of all matching AGO:siRNA complexes) required to increase methylation by 1% was established based on the average dwell time across all loci, assuming that the average dwell time was sufficient to achieve CHH methylation maintenance. For example, as the median methylation was 8% prior to replication, the average dwell time at loci beginning at 8% methylation was sufficient to increase methylation by 4%, resulting in stable maintenance.

### Assessment of solutions

To assess the stability of these simulations, the methylation distribution at the final (10th) generation was compared to the 1st generation. Differences between starting and ending methylation distribution across simulated loci were compared using the non-parametric Kolmogorov-Smirnov test (kstest) in Matlab with a significance threshold of p<0.01. The ‘success’ condition for maintenance of methylation is then p-values greater than 0.01, indicating the final methylation distribution is not significantly different from the starting distribution. Due to the stochastic nature of these simulations, simulation replicates might vary in whether they are scored as non-significantly different, and therefore stable. Simulation conditions that had more non-significantly different final distributions out of 7 simulation replicates were considered more stable. Since the DNA-binding condition requires approximately 3 times longer to run than RNA-binding, only a reasonable fraction of RNA conditions were sampled (**Table 2**).

**Table 2.**
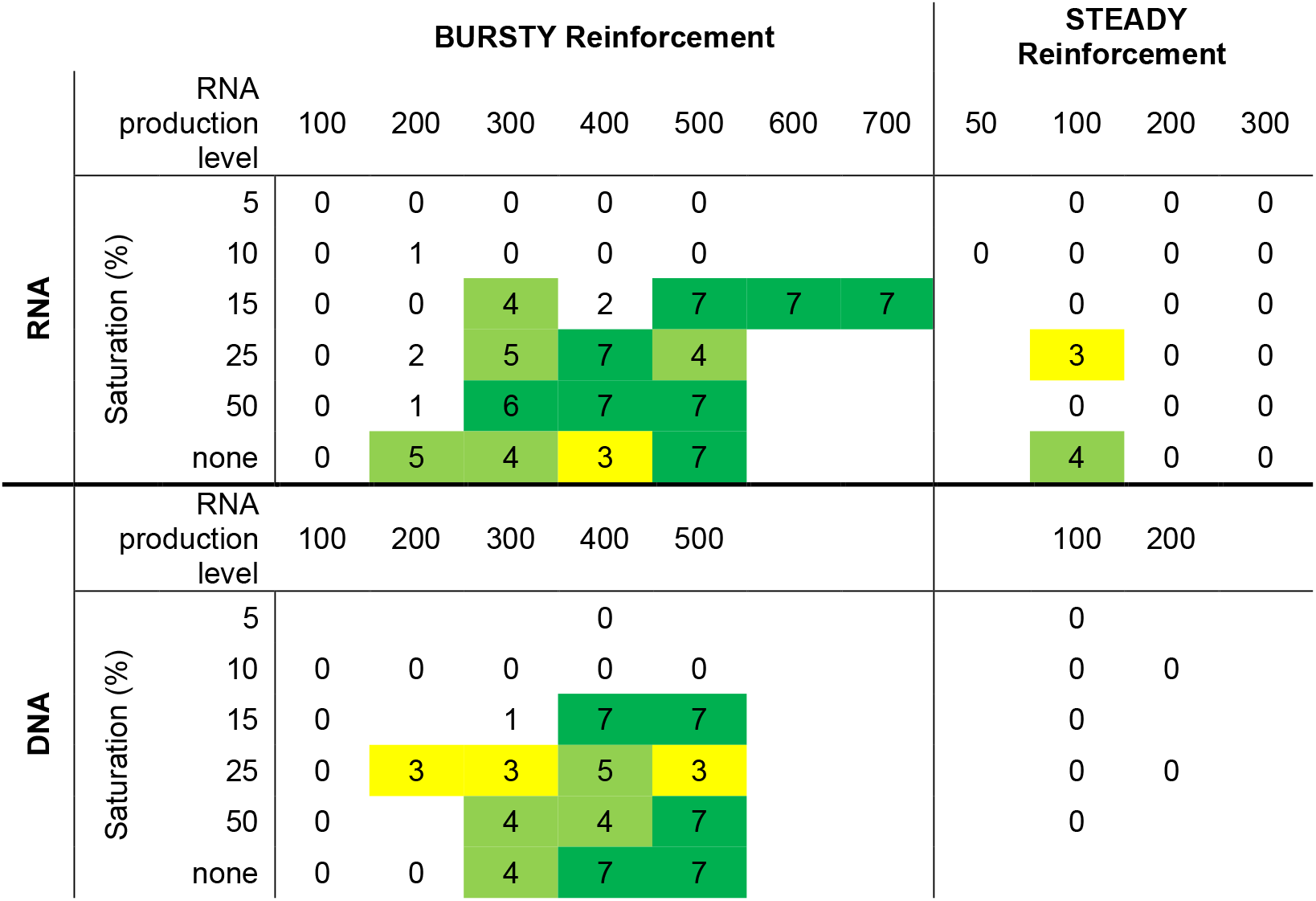
Summary of simulation results. Number of simulations (out of seven) that are insignificantly different from the starting distribution after 10 cycles.

## Results

### A quantitative model for maintenance of DNA methylation via RdDM

Numerous components of RdDM have been identified and the basic mechanism is well understood [4,5]. When constructing our model, we therefore simplified the process to the most critical components, namely siRNA production, siRNA:AGO association with target molecules, induction of methylation, and DNA replication (**Figure 1**).

**Figure 1.**
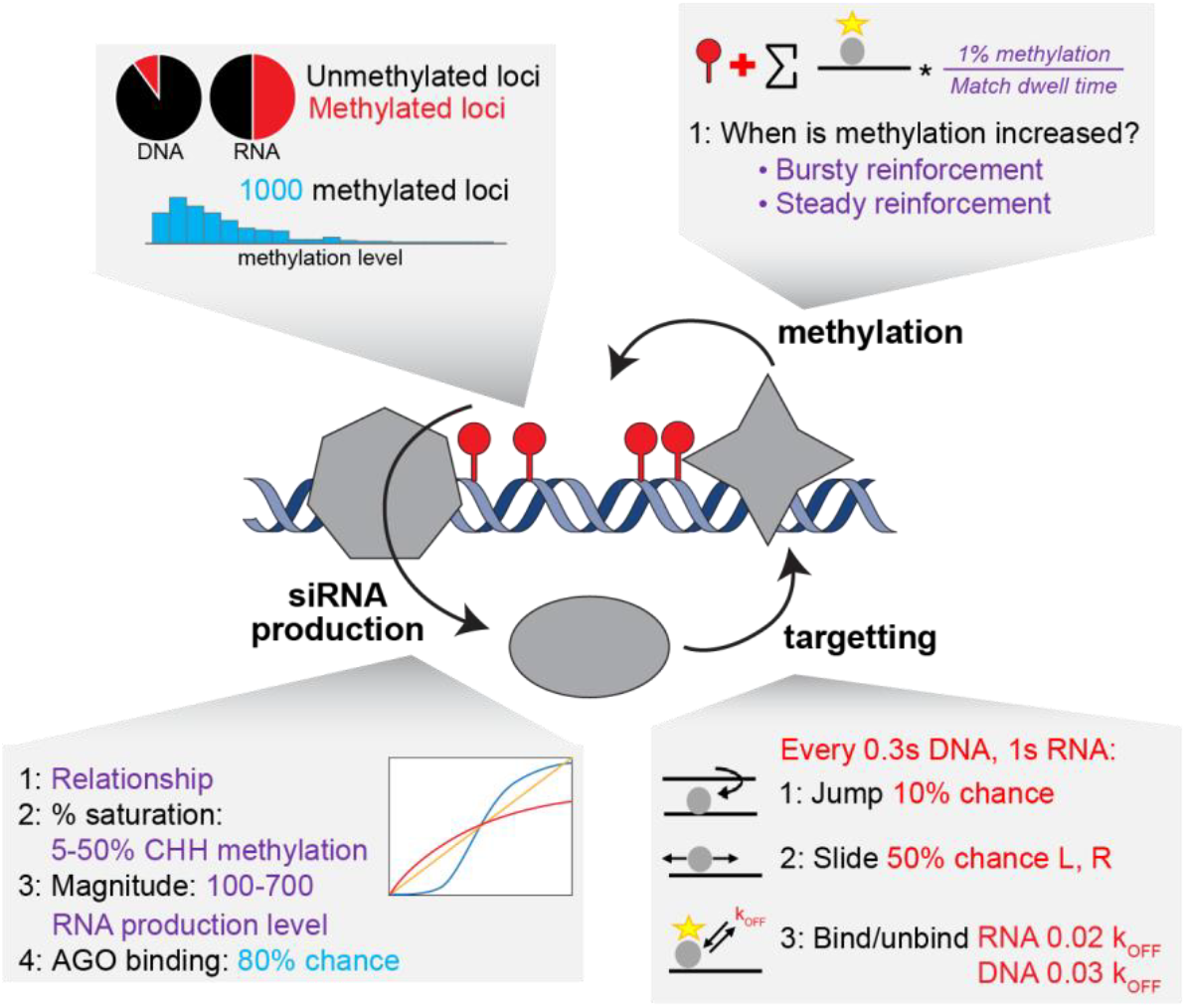
Schematic of CHH model system. Maintenance of DNA methylation by siRNAs is a self-reinforcing loop. SiRNAs are produced by RNA Pol IV and RDR2 (grey heptagon); these siRNAs integrate into AGO proteins (grey oval) and use the sequence of the siRNA to bind target DNA; successful association of the AGO:siRNA with a DNA locus recruits a DNA methyltransferase (grey star) to induce methylation, which causes additional siRNA production. Higher methylation causes greater siRNA production and subsequently more DNA methylation (left), but the process is also stable at lower methylation levels (right). Boxes illustrate the parameters used to model different parts of the system, with purple text indicating values explored in our simulations, blue text indicates fixed values, and red text indicates fixed values obtained from the literature (see **Table 1**).

While 24-nt siRNAs engaged in RdDM are the most abundant class of small RNAs in most plant tissues, the number of siRNAs produced from a given locus per cell is unknown. Similarly, it is not clear whether all siRNAs are bound by AGO4, although there is some evidence to suggest this. AGO4 protein does not accumulate in the absence of siRNAs [12], suggesting that siRNAs might be limiting *in vivo*. However, exogenous siRNAs delivered to mammalian cell culture compete for AGO binding [53], indicating that AGO levels can be limiting in some circumstances. Limiting AGO would primarily influence the ability of loci with low CHH methylation levels to be represented in the AGO:siRNA pool. In preliminary work, we found that the influence of limiting AGO association was easily confounded by changing siRNA production levels, suggesting that the ratio of AGO:siRNA complexes to target molecules is a more important parameter than stochasticity derived from competition between siRNAs for AGO binding. In the simulations presented here, we slightly limit AGO levels (0.8 AGO to 1 siRNA), and compare model behavior across a range of siRNA production levels (**Table 2**).

### Bursts of siRNA production across the cell cycle is favored over constant production

One outstanding question regarding RdDM is when during the cell cycle siRNA production and DNA methylation occur. In fission yeast, small RNA-directed chromatin modification occurs immediately following DNA synthesis, perhaps because newly-synthesized histones lack heterochromatic modifications and therefore are relatively permissive for Pol II transcription [56,57]. However, RdDM is initiated by RNA Pol IV transcription, which is enhanced, rather than repressed, by silent heterochromatin marks [22,58]. To explore the kinetics of siRNA production during the cell cycle, we compared two potential scenarios: “bursty” production, where siRNAs are added to the siRNA pool based on a locus’ methylation level in a single burst immediately after replication, and steady production, which was modeled by siRNA production at six timepoints spread over the course of the cell cycle (*e*.*g*., every 4 hours). In both scenarios, we assume CHH methylation is continuously updated throughout the cell cycle. We compared the performance of these two scenarios across a range of siRNA production levels (100, 200, 300), using a simple linear relationship between methylation level and siRNA production.

After simulating for 10 cycles, we quantified stability by a statistical comparison of the starting methylation distribution and the methylation distribution after the tenth cell cycle. Variation of individual loci might reflect true cell-to-cell variation in methylation that is averaged when a tissue is measured. We therefore focused on whether the final methylation distribution was centered around zero (no change). Each stochastic simulation was simulated seven times. We found that the bursty reinforcement scenario resulted in stable methylation distributions, particularly at higher siRNA production levels. The steady reinforcement scenario performed better at lower siRNA production levels, but remained inferior to bursty (**Figure 2, Table 2**). Final methylation distributions under steady reinforcement demonstrate that some loci become hypermethylated as a consequence of the positive feedback between siRNA production and DNA methylation. We therefore conclude that siRNA production might be limited to a single point within the cell cycle, although this need not be immediately following DNA synthesis, as modeled here.

**Figure 2.**
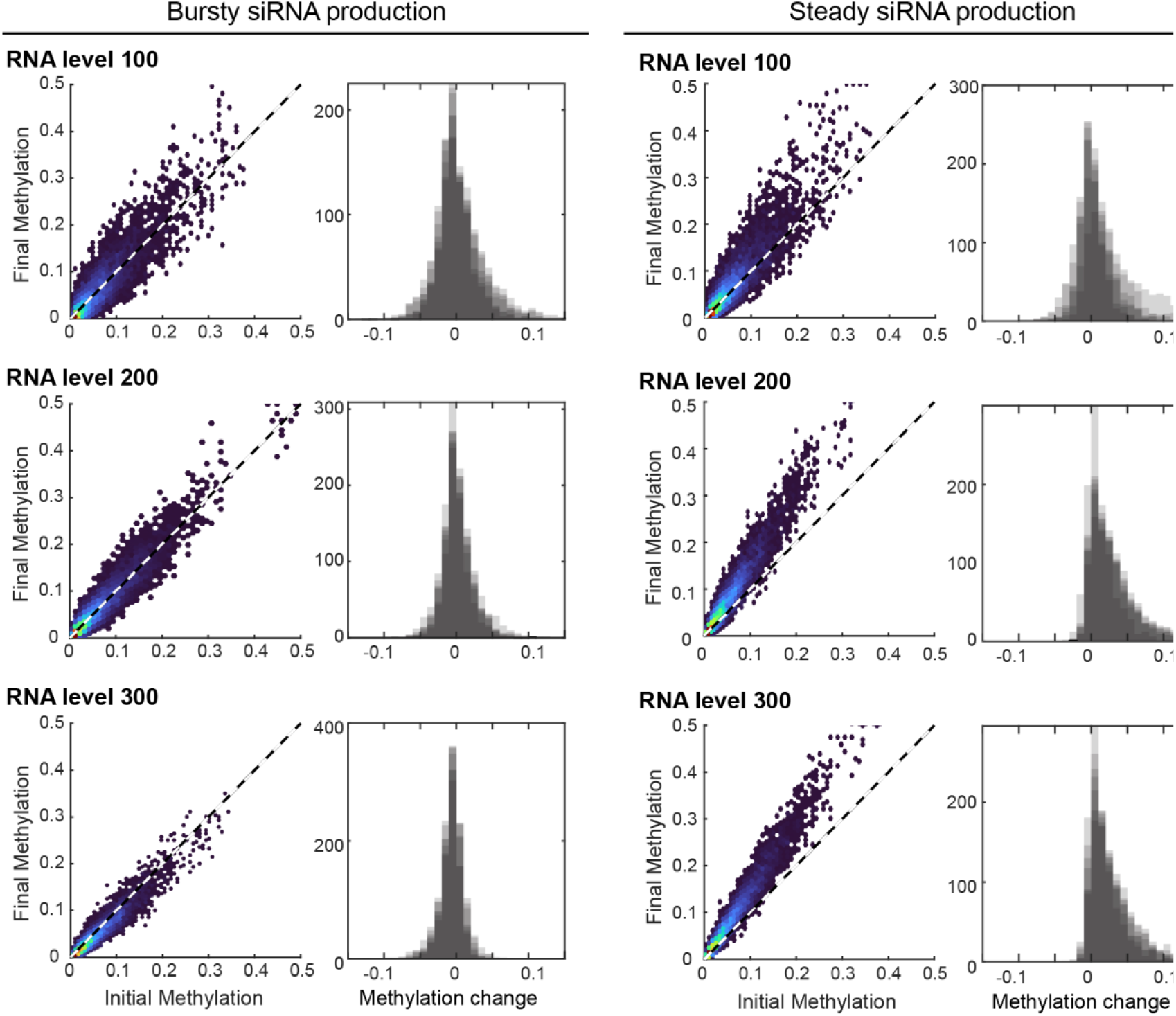
Bursty siRNA production results in enhanced stability of CHH methylation distribution. Varying levels of siRNA production were simulated across ten rounds of DNA replication over seven simulation replicates (all linearly proportional to methylation level) for bursty and steady conditions. Final methylation is plotted versus initial methylation for each locus in a representative simulation (left) and the distribution of methylation change is reported for all seven simulation replicates (right, shaded replicates overlap each other). In bursty conditions, final methylation approximates initial methylation, and changes in methylation are centered at zero. In contrast, steady reinforcement of methylation results in increased methylation relative to the starting distribution. In all cases, higher levels of siRNA production reduced the variance in CHH methylation. The dashed line represents 1:1 correspondence.

### Linear relationships between CHH methylation and siRNA production stabilize RdDM loci with low methylation

Another unanswered question regarding RdDM is the quantitative relationship between DNA methylation and siRNA production. DNA methylation triggers methylation of Histone H3 on Lysine 9 (H3K9me), which in turn recruits siRNA production machinery [22,58,59]. This connection suggests an underlying increasing relationship between siRNA production and CHH methylation. Here, we consider four models: a linear relationship; a linear relationship with a maximum siRNA production, or saturation point, that occurs at a level of methylation (here, we modeled saturation at 5, 10, 15, or 50% methylation); a Hill function relationship; and a logarithmic relationship (**Supplemental Figure S1)**. We ran the model with each of these relationships across a range of siRNA production levels with both bursty and steady siRNA production and quantified the stability of the methylation distribution (**Figure 3, Supplemental Figure S2**).

**Figure 3:**
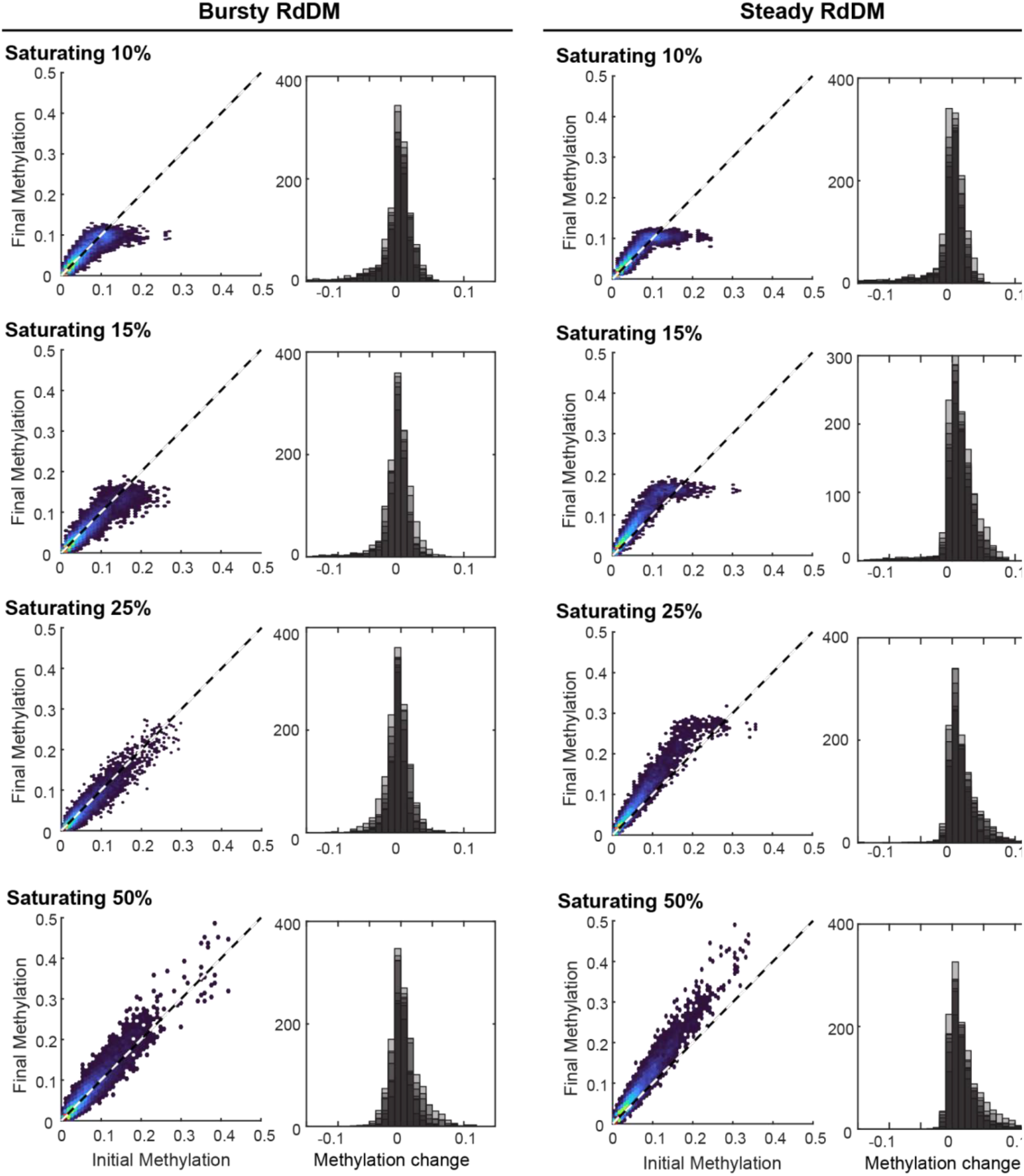
A linear relationship between CHH methylation and siRNA production with a high saturation point is optimal for stable maintenance of CHH methylation. Bursty versus steady reinforcement of methylation was tested with varying levels of saturation (all at RNA factor 300). The saturation level reflects the percent of CHH methylation at which there is no corresponding increase in siRNA production (*i*.*e*., for Saturating 10%, CHH methylation levels of 10% or greater will produce the same number of siRNA). Final methylation is plotted versus initial methylation for each locus in a representative simulation (left) and the distribution of methylation change is reported for all seven simulations (right, shaded replicates overlap each other). In both bursty and steady conditions, saturation results in a plateau of final methylation, however bursty reinforcement with 15% or greater saturation level produces stable maintenance of CHH methylation across 10 rounds of DNA replication and 7 simulation replicates. Results based on non-linear relationships are in **Supplemental Figure S2**.

We found that regardless of bursty or steady methylation reinforcement, a linear relationship between siRNA production and CHH methylation performed the best, with saturation at 15% or higher CHH methylation (**Figure 3**). Note that median CHH methylation is at 8% methylation in our simulations, with 94.8% of loci being less than 15% CHH methylated, and 84.4% being less than 10% CHH methylated. Hill function and logarithmic relationships failed to maintain the initial CHH methylation distribution due to their low coverage of loci with low methylation (**Supplemental Figure S1-S2**).

We also explored high siRNA production levels (500-700) with bursty reinforcement, where siRNA production saturates at 15% CHH methylation (**Supplemental Figure S3**). We found that increasing the siRNA production level under these simulation conditions resulted in stable CHH methylation. Similarly, in models of post-transcriptional RNA silencing, high degrees of stimulus produce more stable behavior [41].

Although RdDM is primarily associated with CHH methylation, there is extensive crosstalk between methyltransferases in plants [60], and the siRNA production machinery might respond to other forms of methylation (*i*.*e*., CG or CHG). We therefore ran the model with the addition of RdDM-independent CG methylation. A level of CG methylation was randomly assigned from a uniform distribution between 0 and 20% methylation and siRNA production was set to be linearly proportional to the total CHH and CG methylation at each locus. The addition of CG methylation resulted in unrealistic CHH methylation distributions that mimic the uniform (0,20%) distribution of CG methylation regardless of RNA production level (see **Supplemental Figure 4**). From these simulations we conclude that the most stable methylation patterns result from siRNA production that is linearly related to the amount of CHH methylation at a locus, and is not meaningfully influenced by the amount of CG methylation.

### Stable methylation is possible with both RNA and DNA target sites

Although most models propose that siRNA:AGO4 complexes associate with non-coding RNA produced by RNA Pol V [14,15,17], it remains possible that these complexes bind to single-stranded DNA denatured by Pol V transcription [16]. Studies of related AGO proteins demonstrate that AGO:siRNA complexes have high affinity for both DNA and RNA strands [44,52,55]. We therefore modified appropriate parameters in the model to test the feasibility of DNA as the AGO:siRNA target and measure systemic differences between DNA and RNA target molecules (**Table 1**). The *k*_*on*_ of AGO to RNA is about 3 times slower than the *k*_*on*_ for DNA [44,52,55]. We therefore used different timesteps when testing AGO:siRNA searching for RNA or DNA targets – every 1 second for RNA and every 0.3 seconds for DNA [44,47]. The *k*_*off*_ was also smaller for RNA than DNA, resulting in shorter dwell times for AGO:siRNA complexes associated with their target DNA site versus a target RNA site. Finally, to account for the greater number of non-RdDM DNA sites in a nucleus, siRNA:AGO complexes were allowed only a 10% chance of encountering one of the RdDM loci in the DNA target scenario, compared to a 50% chance in the RNA target scenario. As the DNA target search simulations take 3 times as long to complete, we limited simulations to those conditions that were observed to be successful or near-successful in RNA binding simulations.

When the model was run with the siRNA:AGO complex binding DNA, we observed that the steady siRNA production condition never produced stable CHH methylation distributions (**Figure 4**). The bursty siRNA production condition maintained CHH methylation, but required a higher siRNA saturation level in comparison to RNA simulation conditions. At higher siRNA production levels (400-500), we found bursty siRNA production resulted in stable CHH methylation distributions at a high rate, similar for both RNA and DNA search conditions. These observations suggest that AGO:siRNA targeting of DNA remains a possible mechanism of RdDM, and would require higher levels of siRNA production.

**Figure 4.**
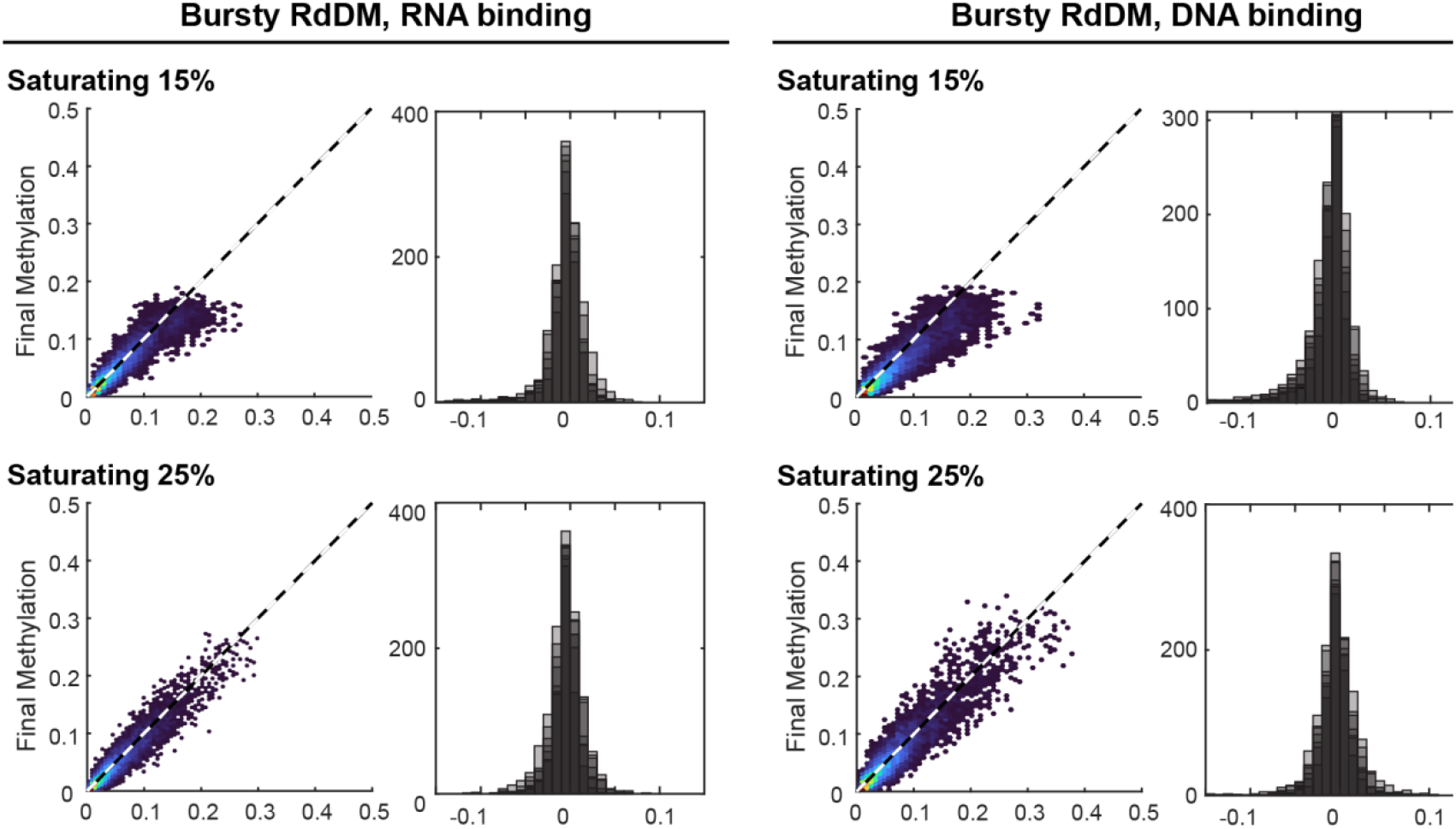
Similar features produce stable CHH methylation maintenance whether AGO searches RNA or DNA. Methylation can be stably maintained bursty siRNA production at two saturation levels when the model is run with either searching RNA or DNA targets. Final methylation is plotted versus initial methylation for each locus in a representative simulation (left) and the distribution of methylation change is reported for all seven simulation (right, shaded replicates overlap each other). See **Table 2** for additional conditions.

### Increasing the level of siRNA production eventually leads to self-inhibition

Although there are clear positive feedbacks in this system, we wondered what prevents the levels of CHH methylation from fully saturating, as in the CG methylation case? We therefore checked the model simulations for predictions of negative feedbacks in our simulations with bursty reinforcement, with a linear relationship between CHH methylation and siRNA production, which saturates at 15% CHH methylation (seven replicates for each siRNA production level). Across these model simulations, as siRNA production increases, the number of AGO:siRNA complexes increase proportionally (**Figure 5**), and the number of loci that lose CHH methylation (CHH methylation reduces to 0%) drops exponentially.

**Figure 5.**
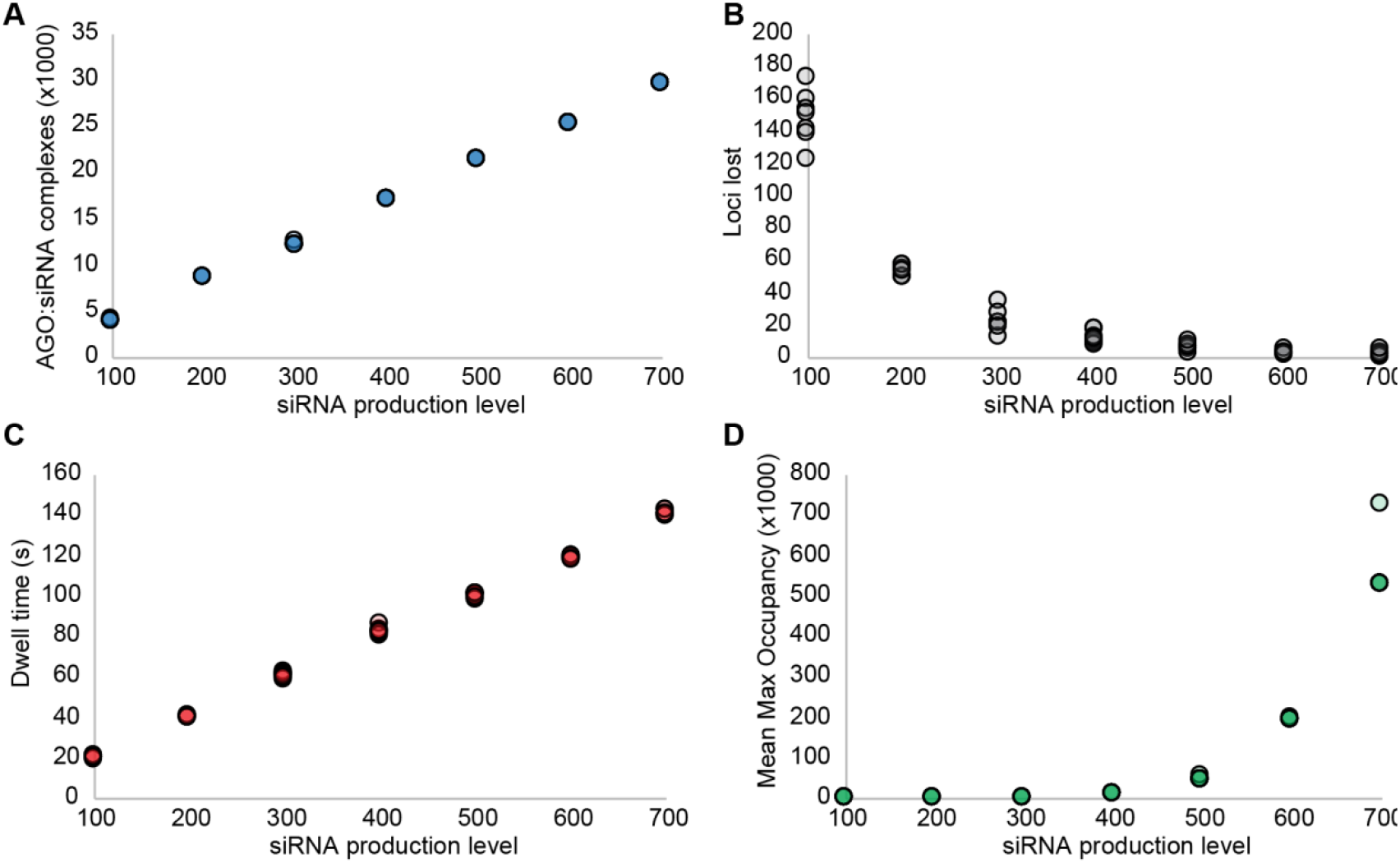
Increasing siRNA production beyond a point leads to self-inhibition. As the level of siRNA production increases, the number of AGO:siRNA complexes increases linearly (A), and the number of loci which lose CHH methylation drops exponentially (B). However, increasing the size of the AGO:siRNA complex pool increases within-complex competition for loci. As the siRNA production level increases, the amount of dwell time (matched AGO:siRNA and loci) to increase CHH methylation by 1% increases linearly (C), and the mean number of times that a methylated loci experiences maximum occupancy increases exponentially (D). In our model, a locus can hold up to 10 AGO:siRNA complexes. Shown is the value of each of 7 simulation replicates for each siRNA production level.

However, the model predicts that increasing siRNA production levels also increases the amount of time to increase CHH methylation by 1%. The amount of dwell time (duration of an AGO:siRNA complex interacting with its matched locus) required to increase CHH methylation by 1% is internally calculated during the first round of every model simulation, as all model parameters influence how long this takes. To calculate this, we assume that the amount of dwell time a locus experiences is sufficient for the median locus to double its CHH methylation fraction (i.e., the amount lost due to DNA replication).

The model also predicts that increasing siRNA production exponentially increases the number of times a methylated locus hits maximum occupancy (10 AGO:siRNA complexes). As we hold the number of CHH methylated loci (1000) constant across all simulations, as the number of AGO:siRNA present in the system increases, eventually there will be more AGO:siRNA available for a given locus than there are areas to interact with that locus.

## Discussion

Here we describe the development of the first mathematical model describing maintenance of CHH methylation by RdDM. We validated the model’s behavior by simulating over a range of parameters and conditions and determined a set of configurations which are able to produce stable maintenance of CHH methylation over a range of starting methylation levels that approximate empirical observations [61]. Although most experimental evidence suggests that CHH methylation due to RdDM is commonly around 20% [30], in this paper we focused on the case of low CHH methylation, with a median of 8%, to determine the conditions required to maintain such an extreme.

Several known features of the CHH methylation system were excluded from our current model for simplicity. For example, we considered siRNA to have locus specificity, but not sublocus site specificity. In reality, an AGO:siRNA complex would only match to a specific site within a locus and might need to slide along the target locus before binding. We also modeled all loci as unique, when many RdDM loci in a genome share homology and siRNAs produced at one locus might be functional at multiple sites. Homologous sites could introduce competition between loci, which might reduce stability of methylation; alternatively, siRNA production at a homologous locus could restore methylation levels that had been lost and thereby buffer the system. Similarly, our model was completely cell autonomous, whereas siRNAs are known to move intercellularly and function non-cell autonomously [62]. Intercellular movement of siRNAs (or siRNA:AGO complexes) also offers the possibility of both competition and mutual support between CHH methylated loci. Most importantly, we assumed that all AGO:siRNA complexes disassociate during DNA replication and must randomly rediscover their target loci. Nothing is known regarding the fate of RNA Pol V or its transcripts during DNA replication and it remains possible that AGO:siRNA complexes might be preserved at their target locus in some manner, perhaps in association with the Pol V carboxy terminal domain. Despite these simplifications, the model provides insight into the quantitative features required for stable maintenance of CHH methylation by RdDM.

Firstly, we find that bursts of siRNA production, wherein the pool of siRNA is filled right after replication, results in more stable CHH methylation distributions under a range of simulation conditions compared to steady reinforcement, where siRNAs are produced throughout the cell cycle. Bursty siRNA reinforcement has been observed at the transcriptionally silent pericentromeric repeats in fission yeast and might explain the paradox of Pol II transcription being required to establish transcriptionally silent chromatin [56,57]. Because DNA synthesis reduces H3K9me, fission yeast pericentromeres become permissive for Pol II transcription allowing the production of siRNAs to reestablish H3K9me. However, such a compensatory dynamic is not expected at RdDM loci in plants, because DNA methylation and H3K9me promote siRNA production rather than inhibit it. A mechanism restricting siRNA production to a single point of the cell cycle is unknown, but might involve other histone modification or the density of linker histone [23]. Regardless of the mechanism, bursts of siRNA production would answer at least one outstanding question: how transcription of the same locus by RNA Pol IV and Pol V is coordinated. It might be that the functions of these polymerases are temporally separated during the cell cycle.

Secondly, our simulation results strongly support a linear relationship between siRNA production and CHH methylation, as alternative biologically-relevant relationships (Hill function and sigmoid function) resulted in loss of methylation distribution. Under a linear relationship, saturation of siRNA production at 15% CHH methylation or higher was also sufficient to maintain CHH methylation. However, very few of the loci in our model existed at methylation levels above this saturation point, and it is it not clear whether higher methylation can be maintained under saturation. The simulations also suggest that the number of siRNA produced per percent CHH methylation needs to be sufficiently high to achieve stable maintenance of CHH methylation.

Although our results favor the linear relationship over the nonlinear relationships explored, it is unclear what biological mechanisms might produce linearity. Due to the computation time required to run detailed simulations, we do not explore every possible relationship between methylation and siRNA production. We interpret our results as not supporting a linear relationship *per se*, but rather a relationship sufficiently linear to satisfy requirements of this system. For example, our results suggest that it is critical to have siRNA coverage of those loci with low levels of CHH methylation, while higher degrees of CHH methylation can be maintained with relatively lower representation in the siRNA pool. Lower production of siRNA for loci with high degrees of CHH methylation would reduce competition for the locus, and for AGO generally, by reducing the total siRNA pool size.

Finally, stable maintenance of CHH methylation was possible whether the model was set for AGO:siRNA searching of RNA or DNA. Although all characterized AGO:sRNA systems are demonstrated or presumed to target RNA, including those that cause transcriptional silencing of chromatin [17,63,64], there is also evidence for AGO4-DNA association during RdDM [16]. Our model demonstrates that targeting of DNA by AGO:siRNA complexes is feasible and also that the viable parameter space for these two targets overlaps, offering the possibility that cells might enable targeting of DNA and RNA simultaneously.

CHH methylated loci have the apparent ability to stably maintain a wide range of methylation fractions, while CG methylated loci tend toward bi-modality, exhibiting saturation or depletion of CG methylation. There is a strong positive feedback loop in the CHH methylation reinforcement pathway, where more AGO:siRNA dwell time increases CHH methylation levels, which produce additional siRNA, enabling more AGO:siRNA complexes, and so on. The model predicts that careful tuning is needed in the negative feedbacks present in this system to stably maintain a range of CHH methylation fractions (**Figure 6**). First, DNA replication is a strong negative feedback, reducing both CHH methylation fractions and the AGO:siRNA complex pool by half. Secondly, bursty reinforcement rather than steady reinforcement limits the degree to which the system can accumulate siRNA and consequently CHH methylation by limiting the strength of that positive feedback. The model also predicts that over-production of siRNA leads to self-inhibition, particularly as the number of AGO:siRNA complexes surpasses the capacity of their matched methylated locus.

**Figure 6.**
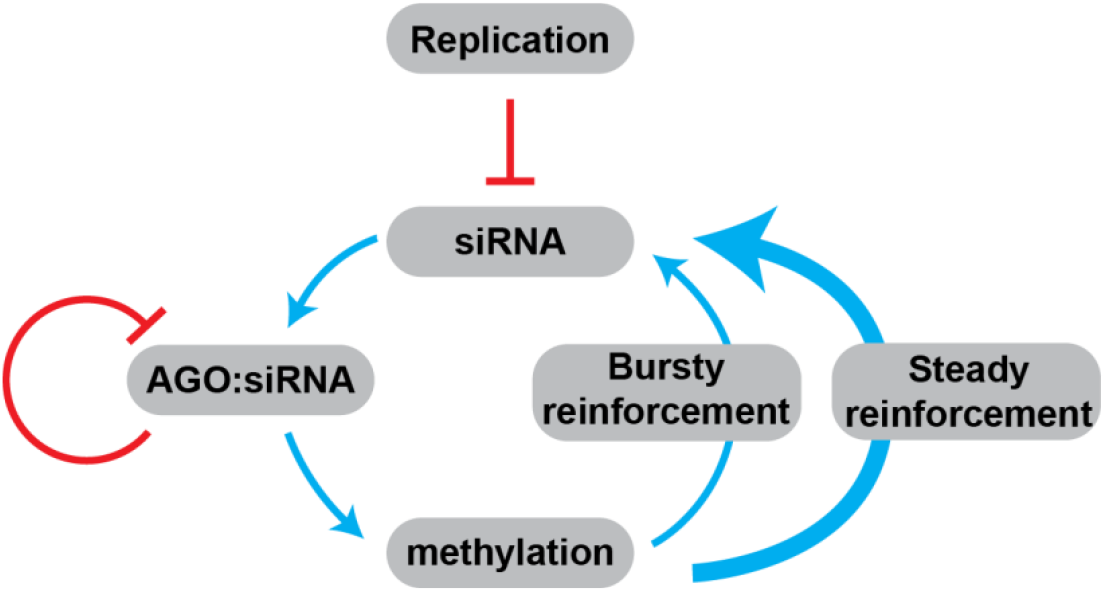
Stable maintenance of a range of CHH methylation fractions requires balance of negative feedbacks with methylation reinforcement. CHH methylation reinforcement produces a strong positive feedback loop, where increasing CHH methylation at a locus increases siRNA production from that locus, and so on. DNA replication provides strong negative feedback on the CHH methylation system, reducing both methylation and siRNA by half. Our modeling results predict that stable maintenance of CHH methylation is enabled by bursty reinforcement, which limits the strength of that positive feedback loop, as well as internal competition between AGO:siRNA complexes for their matched loci.

Epigenetic pathways like RdDM are inherently difficult to understand due to the nature of their self-reinforcing states - once one aspect is disrupted, the entire system collapses. For many years, geneticists have used this fact to identify components of RdDM and biochemists have then investigated the physical interactions and enzymatic activities of these components in isolation. Our mathematical model demonstrates the insight to be gained by applying quantitative modeling to epigenetic systems. We hope it will serve as a launching point for additional research in this area.

## Author Contributions

R.D. and R.A.M. conceived and developed the project; R.D. created the model and ran all simulations; R.D. and R.A.M. wrote the manuscript.

## Acknowledgements

This work was supported by the National Science Foundation (IOS-2109790 to R.D., IOS-1546825 and IOS-2247914 to R.A.M.) and a Find Your Inner Modeller (FYIM IV) Travel and Training Award (NSF MCB-2003415). The authors also thank Dr. Tania Chakraborty for discussion and data sharing.

