## Supplemental Figures for "Mathematical model of RNA-directed DNA methylation predicts tuning of negative feedbacks required for stable maintenance"

### Dale and Mosher, Supplemental Online Material

#### Supplemental Text 1. Model Algorithm

Model algorithm

1. **Simulation initial conditions.** Simulator conditions that are set in the model include the relationship between siRNA production and methylation level; the type of relationship between siRNA production and methylation level (*i.e.*, linear, sigmoid, Hill function); whether the relationship between siRNA production and methylation level becomes saturated at a given methylation level (*e.g.*, 10% methylation, 25% methylation); whether or not CG methylation contributes to siRNA production; if the AGO:siRNA complex searches RNA or DNA; the initial methylation distribution across the loci; the number of loci; the number of days to be simulated; and whether reinforcement occurs in a steady or bursty fashion.
2. **Creating the loci.** Each of *N* loci has 12 segments – 10 binding segments and a 0th and 11th segment which represent spatially near sites that are off-locus. These numbers were chosen to simulate a 200 nucleotide long siRNA-producing locus, with 20 nucleotide long siRNA. An AGO:siRNA complex is considered on a matched locus if it is on segments 1-10. A locus is considered saturated when 10 AGO:siRNA complexes are present, and additional AGO:siRNA complexes are unable to interact. Each siRNA has locus and segment specificity, but in these simulations, segments are otherwise not tracked, and multiple AGO:siRNA complexes are allowed to co-occupy a given segment as long as the total number of AGO:siRNA complexes on a locus does not exceed the threshold.
3. **Assigning methylation levels.** The initial methylation distribution is set using the distribution $-log(1-(1-exp(-\mu)) * U \;/\; \mu$ , where $\mu$ is the mean methylation fraction, and $U$ is uniformly-distributed noise, $1 - .9 * Unif(0,1)$. This distribution was chosen as it can take a variety of forms (left or right skewed, symmetric, and uniformly distributed). The default case takes a value of $\mu$=10, which produces a median CHH methylation level of 8% prior to replication, or 4% CHH methylation post-replication. For *N* methylated loci, *N* random CHH methylation fractions are drawn from this distribution. Immediately after assigning methylation levels, the levels are cut in half to represent loss due to DNA replication.
4. **Creating AGO:siRNA pool:**
    a. Each methylated locus has a fraction of CHH methylation. The number of steady-state siRNAs each locus is able to produce is determined by the siRNA-to-methylation relationship (*i.e.*, linear, sigmoid). The number of siRNA this locus produces is then calculated. In the linear case, the linear parameter (*e.g.*, 1) would be multiplied by the CHH methylation level (*e.g.*, 8%) and the siRNA production parameter (*e.g.*, 100). This results in a pool of siRNA from each locus relative to its methylation level (*e.g.*, 8 siRNA). Each siRNA has locus and segment specificity.
    b. If AGO is limiting as a fraction ***f*** (*e.g.*, .8 AGO to 1 siRNA), then each siRNA from each locus has a chance of ***f*** to enter the AGO:siRNA pool. If AGO is limiting as

an objective total poolsize, then the resulting fraction ***fr*** is calculated, and each siRNA has an ***fr*** chance of entering the pool. It should be noted that changes in CHH methylation will therefore result in changes in AGO:siRNA complex poolsize, further exacerbating departures from the initial CHH methylation distribution.

5. **DNA target search.** AGO:siRNA randomly interacts with a random loci out of all possible *N* loci, including non-methylated loci. There is a 50% chance to interact with a non-methylated locus if an AGO:siRNA complex searches RNA, or a 90% chance if searching DNA. Upon interaction with a locus, the AGO:siRNA complex is given a random segment number (0-12).
    a. If a locus is full (10 AGO:siRNA are present), the AGO:siRNA complex instead interacts with another random locus.
    b. Once on a locus, it has a 45% chance to slide left, 45% chance to slide right, and 10% chance to leave the locus.
    c. If it slides off-locus (*i.e.*, the -1st or 12th segment by sliding left or right), or if it leaves the locus, it is assigned a new random (with replacement) locus from all possible loci.
    d. If the AGO:siRNA complex is on its matching locus, it is considered interacting with that locus. The time-to-dissociate is determined by its $k_D$, using the waiting time distribution (Exponentially distributed) calculation in time step increments. The simulation time step is determined by AGO:siRNA complex's $k_{on}$, corresponding to 0.1 seconds when searching RNA, and 0.3 seconds when searching DNA. The waiting time distribution given amount of time on a locus results in a chance-to-leave. If a coin flip parameter alpha is greater than this chance, the AGO:siRNA complex remains on the locus. If the AGO:siRNA complex leaves the locus, it is randomly assigned a new locus with replacement. Every timestep that a locus has a matching AGO:siRNA complex is summed to calculate the total dwell time for that locus, representing that locus' capacity to attract the DNA methyltransferase DRM2.
6. **Connecting AGO:siRNA complex matching to methylation.** After the first simulation day ('burn-in' day), the mean dwell time of the median methylation level is calculated. This mean dwell time parameter determines the dwell time required to maintain methylation. For example, if the median CHH methylation level is 8% pre-replication (4% post-replication), and the corresponding mean total dwell time is 100 seconds, then under stable conditions 100 seconds of total dwell time is required for a locus to increase its methylation by 4% (recoup the reduction due to replication). Thus, in this example, 25 seconds of total dwell time produces an increase in CHH methylation of 1% across all loci in the simulation for the remaining simulation days.
7. **Replication and reinforcement.** After the first day, replication occurs, and the CHH methylation level of all loci are cut in half. The AGO:siRNA complex pool is randomly reduced by half. After the burn-in day, the AGO:siRNA complex pool can be reinforced.
    a. In bursty conditions, reinforcement occurs immediately following replication. The siRNA levels are updated by comparing the maximum level of siRNA in the pool (*e.g.*, *relationship (RNA factor * methylation level)*) to the actual level in the

retained pool. This provides the new siRNA pool, which enters the AGO:siRNA complex pool as in step 4b, subject to any limiting AGO conditions.

b. In steady conditions, reinforcement occurs every 4 hours (6 times a day), chosen as a reasonable frequency. Every 4 hours, loci methylation levels are updated, current and maximum siRNA levels are compared to create a new siRNA pool, which enters the AGO:siRNA complex pool as in step 4b, subject to any limiting AGO conditions.

8. **When CHH methylation is lost.** Loci fail to produce siRNA when they reach *relationship (RNA factor * methylation %)* <1. Loci are considered lost when methylation is below this value, however such a locus may have some siRNA in the retained pool, and are able to recover.

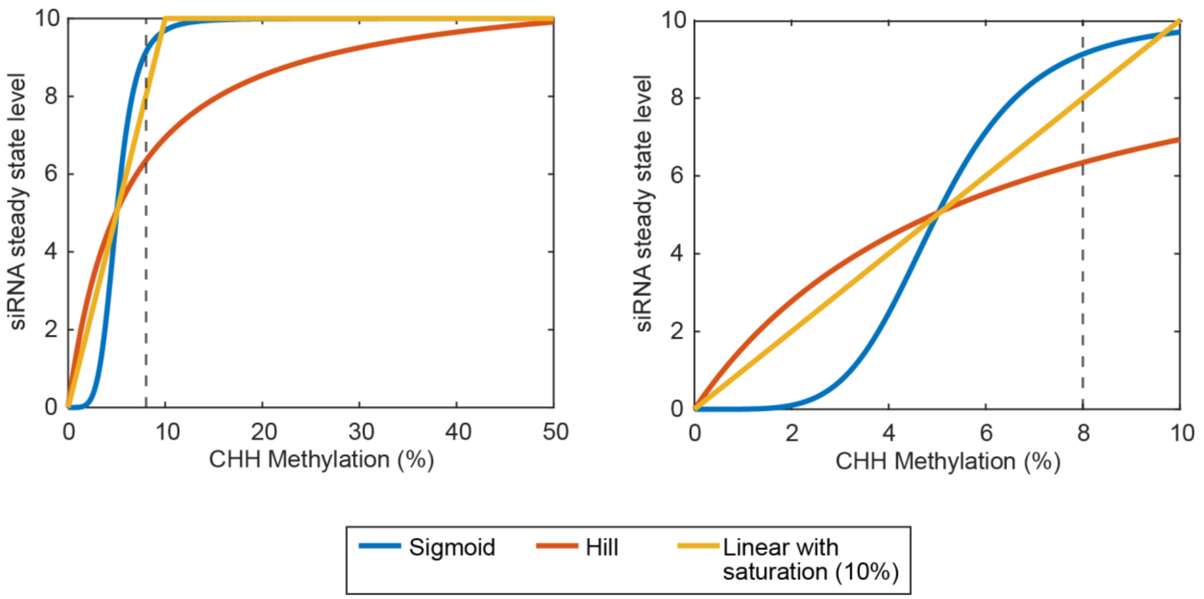


**Supplemental Figure S1. Non-linear relationships between CHH methylation and siRNA production.** We compare the steady-state siRNA production level with the fraction of CHH methylation under linear and non-linear (Hill function and sigmoidal) siRNA production models across the entire range of CHH methylation level (left) and zoomed in (right). The median CHH methylation level in our simulations is 8%. Compared to the linear model, the sigmoid function model has poor coverage in the very low CHH methylation range, while the Hill function model has low coverage in the high CHH methylation range.

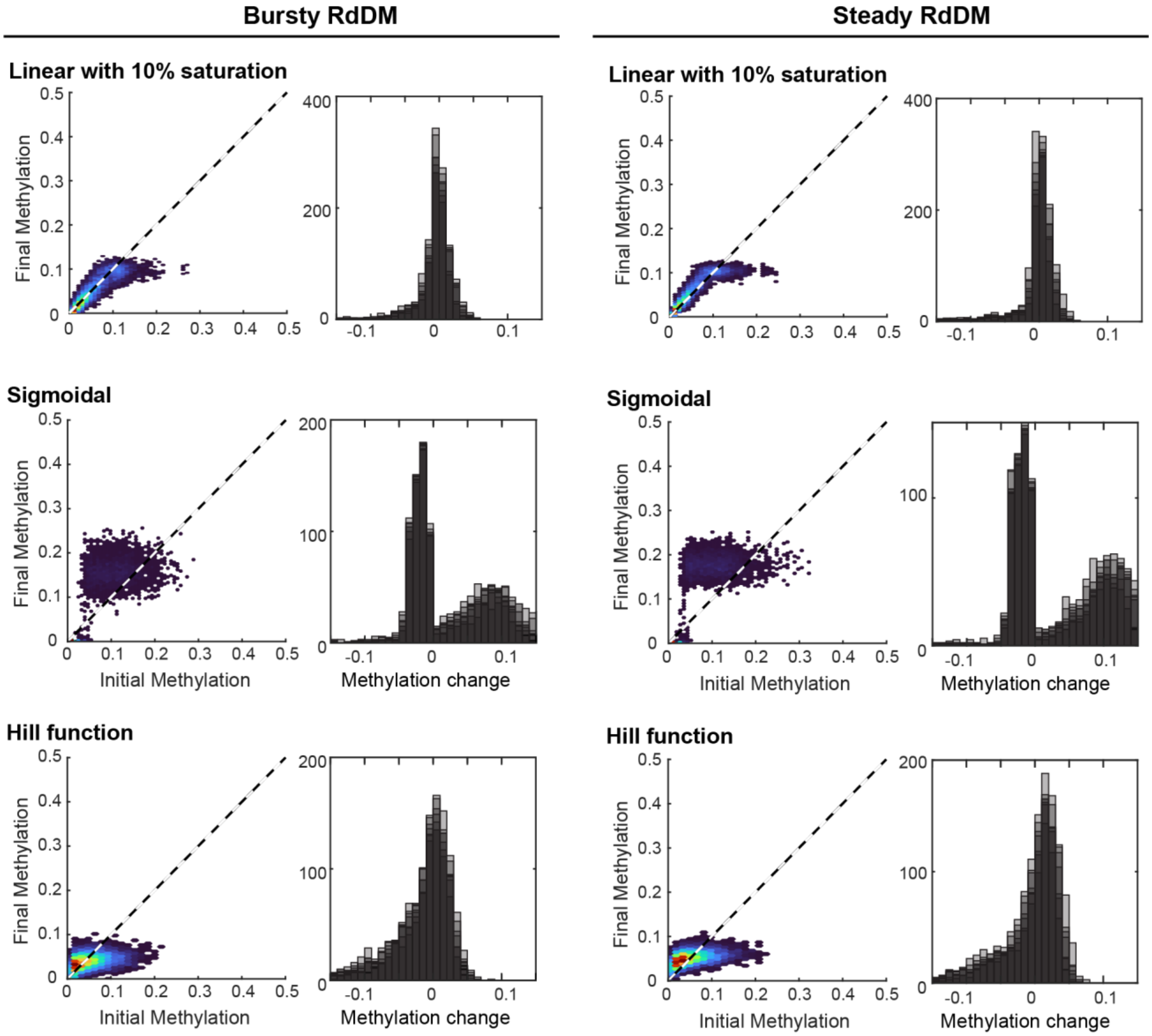


**Supplemental Figure S2. Non-linear relationships between CHH methylation and siRNA production fail to maintain stable maintenance of CHH methylation due to low coverage.** Hex plots comparing starting and final methylation for one representative simulation and histograms of methylation change for seven replicate simulations (overlapped) for Hill function and sigmoidal relationship. Linear with saturation plots are the same as in Figure 3. For both bursty (left) and steady (right) siRNA production, the linear model performs the best. The sigmoidal model causes an increase in CHH methylation, while the change in CHH methylation is bimodal, with many loci either being lost or increasing. The Hill function model causes a reduction in CHH methylation level, and the change in CHH methylation is skewed negative.

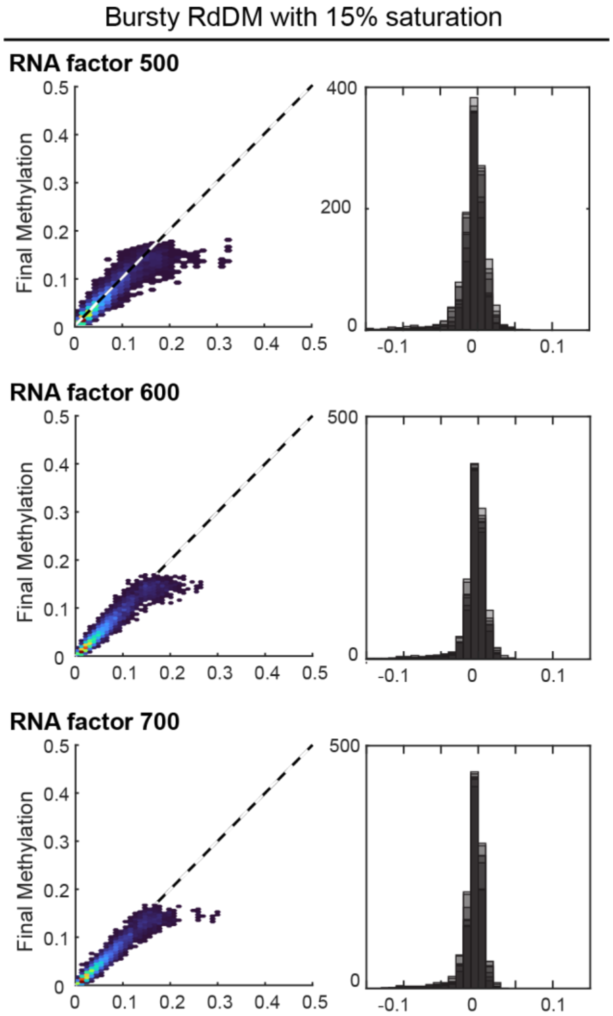


**Supplemental Figure S3: Increasingly high levels of siRNA production improve CHH methylation stability.**
Bursty RdDM with 15% saturation more closely maintains methylation patterns at very high siRNA production levels (RNA factor 600, 700). Final methylation is plotted versus initial methylation for each locus in a representative simulation (left) and the distribution of methylation change is reported for all seven simulation (right, replicates overlap each other).

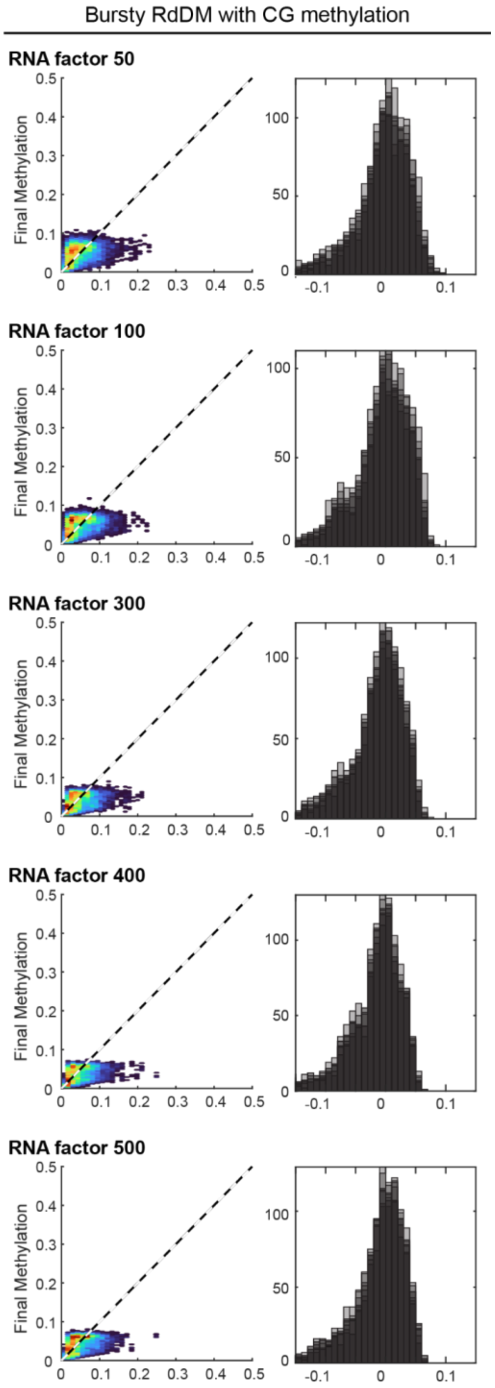


**Supplemental Figure S4. The model suggests that contribution of CG methylation to RdDM results in suppressed CHH methylation levels.** Here we simulated the inclusion of CG methylation (random uniform distributed from 0-20% across CHH methylated loci) in the total methylation level involved in RdDM (*i.e.*, CHH + CG methylation is proportional to siRNA production level). Across a wide range of siRNA production levels (here, RNA factor) we see a suppression of CHH methylation, resembling a uniform distribution between 0-10%, resembling the uniformly distributed CG methylation (between 0-20% prior to DNA replication corresponds to 0-10% post-replication, which is shown in the figures).
